# MINA: Do frozen DNA encoders beat sequence composition? A homology-aware linear-probe benchmark

**DOI:** 10.64898/2026.05.25.727711

**Authors:** Austin Senna Wijaya, Hayden Leung

**Affiliations:** Department of Computer Science, Columbia University, New York, NY 10027, USA

## Abstract

Frozen DNA encoders are widely used as feature extractors, but fine-tuning hides what their fixed embeddings already make linearly accessible. MINA is a lightweight benchmark that asks whether those embeddings beat sequence composition, with two linear probes on 3,244 human genes: 5-way classification into coarse protein families, and regression onto each gene’s GenePT text-summary embedding. Mostly, they do not. On coding sequence (CDS) under a homology-aware split, the validation-selected encoder, NT-v2, recovers coarse family signal at about the level of nucleotide composition (macro-F1 0.693 vs. 0.639; Holm *p* = 0.108) but does not reach amino-acid composition (0.767) or the protein language model ESM-2 (0.954). On GenePT, DNABERT-2 beats nucleotide composition (*R*^2^ 0.065 vs. 0.039) but not amino-acid composition (0.080). Evaluation choices move scores as much as the encoder does: pooling alone takes NT-v2 from 0.504, below nucleotide composition, to 0.698; a mostly noncoding transcription-start-site window, where any signal is likely regulatory rather than family, drops NT-v2 from 0.694 to 0.300 on the same genes; and a random split infiates it from 0.693 to 0.801 through homologous genes leaking into the test set. Frozen DNA encoders thus recover little from coding sequence beyond nucleotide composition, and fair comparisons must use composition baselines and fix pooling, substrate, and split.

**Availability:** code, splits, and artifacts: github.com/Austin-Senna/dna_to_text; sequences, features and predictions: Zenodo, doi:10.5281/zenodo.23031890.

## 1 Introduction

Pretrained DNA encoders such as DNABERT-2 (Zhou et al., 2024), Nucleotide Transformer (Dalla-Torre et al., 2025), GENA-LM (Fishman et al., 2025), and HyenaDNA (Nguyen et al., 2023) are increasingly used as genomic feature extractors, yet their embeddings, despite differing tokenisation, architecture, and pretraining corpus, are often treated as interchangeable summaries of biological sequence. Strong downstream performance after fine-tuning does not reveal what the encoder already represents; linear evaluation (Zhang et al., 2016; Alain and Bengio, 2016), training only a linear readout (a *probe*) on frozen features, asks that narrower question.

We ask two questions of frozen DNA embeddings: do they carry protein-family signal beyond what sequence composition already provides, and do they carry gene function as described in text? To answer them we introduce MINA (Model Interrogation of Nucleotide Architectures), a lightweight linear-evaluation benchmark for frozen DNA encoders on 3,244 human protein-coding genes (Figure 1). One probe classifies each gene into one of five broad protein families; the other regresses its embedding onto the gene’s GenePT vector, a 1,536-d OpenAI text embedding of the gene’s short NCBI summary of what it does (Chen and Zou, 2024). Both read from two substrates: the coding sequence (CDS), which carries protein-domain signal directly, and a long transcription-start-site (TSS) window, which is mostly noncoding.

**Figure 1:**
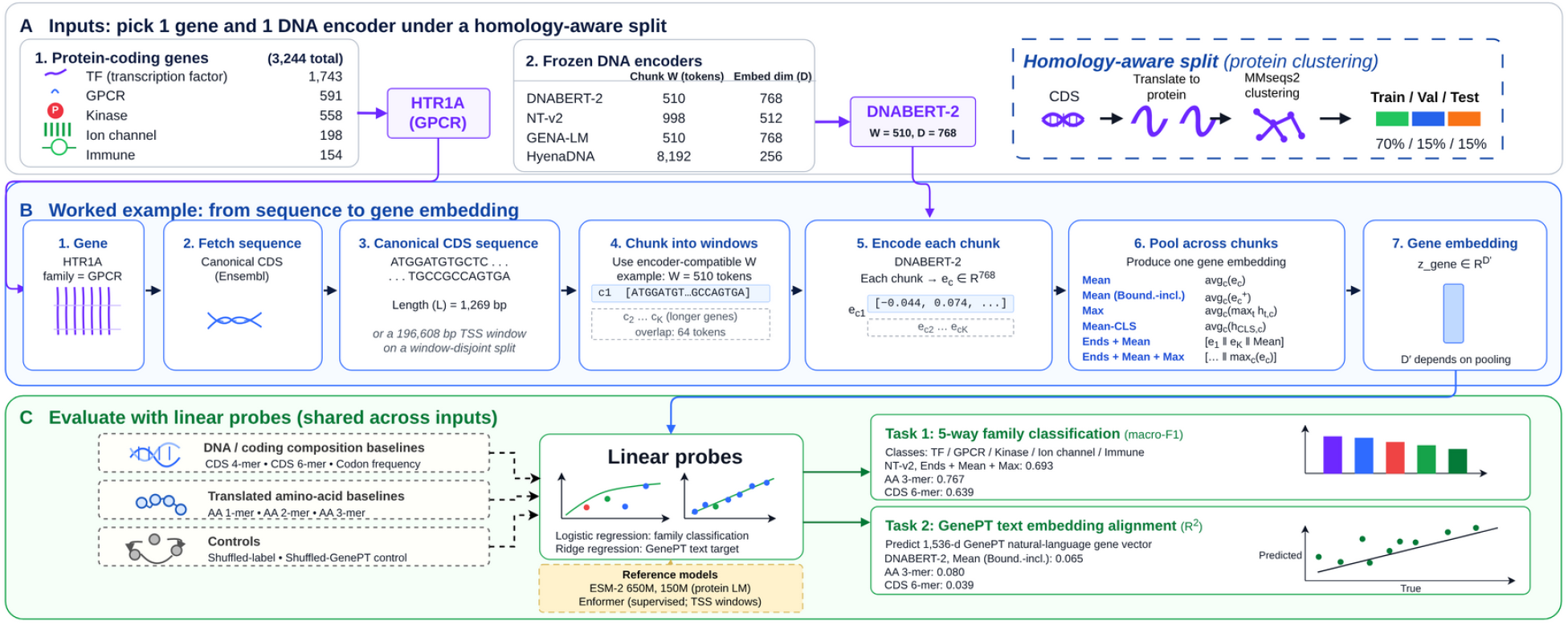
Study design. (A) Genes, encoders (W: content tokens per chunk), and the homology-aware split. (B) A gene’s CDS is chunked, encoded, and pooled. (C) Linear probes against composition baselines, controls, ESM-2, and Enformer; printed scores are CDS test values for the encoder, pooling, and k chosen on validation.

Answering these questions fairly requires three controls. Frozen DNA-encoder embeddings probed on regulatory tasks offer little advantage over conventional models trained on one-hot sequence (Tang et al., 2025), so every probe is anchored by nucleotide and translated amino-acid composition, with ESM-2 (Lin et al., 2023) as a protein-level upper bound and a supervised En-former comparator (Avsec et al., 2021) on TSS windows. Homology between train and test inflates genome-trained models (Rafi et al., 2025), echoing homology-aware splitting in protein machine learning (Li and Godzik, 2006; Rost, 1999; Teufel et al., 2023), so genes are split by protein similarity with no close homologs shared between training and test. The pooling rule that turns chunk embeddings into one gene vector, usually left at a default, is varied across six rules.

Existing DNA-LM benchmarks do not combine these controls. GUANinE (Robson and Ioannidis, 2024) and BEND (Marin et al., 2024) evaluate DNA models on regulatory and genome-annotation tasks. Neither asks what DNA embeddings of coding sequence carry about the encoded protein; BEND restricts variant effects to noncoding regions because coding effects can be predicted from the protein. DART-Eval (Patel et al., 2024) finds that on regulatory DNA, DNA language models do not clearly beat simpler baselines; MINA extends that test to coding sequence, with composition and protein-level anchors. GUE (Zhou et al., 2024) ranks encoders after fine-tuning, with no non-LM baselines. None anchors DNA encoders against amino-acid composition and a protein language model, the comparison MINA is built around.

Our contributions are threefold:

(i) MINA, a linear-evaluation benchmark for frozen DNA encoders, anchored by composition and a protein language model under a homology-aware split;
(ii) evidence that, under linear probes, frozen DNA encoders recover coarse protein-family signal from coding sequence at about the level of nucleotide composition, do not reach amino-acid composition, and recover little text-described function;
(iii) a quantification of how pooling, substrate, and train/test split each change a benchmark’s conclusion.

## 2 Methods

MINA fixes a task and varies how it is evaluated. The task (§2.1) is a pair of linear probes on frozen DNA embeddings of 3,244 human genes, read against composition and protein-level anchors, as shown in Figure 1. The evaluation axes (§2.2) are the pooling rule (Table 1), the input substrate, and the train/test split.

**Table 1:** Pooling rules. Each reduces an encoder’s chunk token embeddings to one gene vector; ∥ is concatenation. CLS is the summary token BERT-style tokenizers prepend to each input (Devlin et al., 2019); HyenaDNA has none.

| **Rule** | **Definition** |
|---|---|
| Mean | mean over tokens in each chunk, then over chunks |
| Mean (Boundary-including) | Mean, including the CLS/SEP boundary tokens |
| Max | max over tokens in each chunk, then mean over chunks |
| Mean-CLS | CLS token of each chunk, mean over chunks |
| Ends + Mean | first chunk ‖ last chunk ‖ Mean |
| Ends + Mean + Max | Ends + Mean ‖ max over chunks |
| TSS-Anchored | the chunk nearest the TSS (TSS windows only) |

### 2.1 The benchmark

#### Genes and targets

We curated 3,244 human protein-coding genes from five functional families: transcription factors (1,743), G-protein-coupled receptors (591), protein kinases (558), ion channels (198), and immune receptors (154). Genes were selected by matching each family’s name against the HUGO Gene Nomenclature Committee (HGNC) gene-group annotations (Seal et al., 2023), intersected with the GenePT v2 gene-symbol set (Chen and Zou, 2024); genes matching more than one family were assigned first-family-wins in the order kinase *→* transcription factor *→* ion channel *→* GPCR *→* immune receptor. Canonical coding sequences (CDS) were retrieved from Ensembl (Harrison et al., 2024). Each gene carries two targets: its family label and its 1,536-d GenePT vector.

#### Encoders

We compared four frozen self-supervised DNA encoders on canonical CDS. Three are BERT-style masked-language models with *k*-mer or byte-pair tokenisation: **DNABERT-2** (Zhou et al., 2024), **Nucleotide Transformer v2 multi-species 100M** (Dalla-Torre et al., 2025) (hereafter NT-v2) and **GENA-LM base** (Fishman et al., 2025). The fourth, **HyenaDNA large** (Nguyen et al., 2023), is an attention-free causal single-nucleotide model. The four span 6.6 M–117 M parameters and 512–2,048-token (up to 1M-nucleotide, for HyenaDNA) context windows; full checkpoints, tokenisation, and architecture details are given in Appendix A.2. All encoder weights were frozen.

#### Probes

Each probe asks what a linear readout recovers from a frozen embedding. For family classification we trained multinomial logistic regression, scored by macro-F1, which weights each family equally, and by Cohen’s *κ* (Vieira et al., 2010), which weights each gene equally after discounting chance agreement. For GenePT we trained ridge regression to predict all 1,536 dimensions of each gene’s GenePT vector, and report *R*^2^ averaged across dimensions, where averaging weighs every dimension equally so a few high-variance dimensions cannot inflate the score. Cosine similarity between predicted and true vectors is a secondary measure. Every choice is made on the validation genes: the regularisation strength for every probe, the composition *k*, and, for the DNA encoders, the pooling rule and the encoder. We then refit on training and validation genes together and score once on the held-out test genes (grids in Appendix D.1).

#### Anchors

Our probes have four reference points: **Composition**: parameter-free *k*-mer counts (Appendix Table A2), which test whether an encoder recovers more than sequence composition. Nucleotide *k*-mer counts are the fioor: like the encoders, they ignore reading frame. On CDS, *k* is selected on validation (*k* = 6 for family classification). In-frame codon counts add the reading frame; amino-acid *k*-mer counts on the translated protein set the higher bar of protein composition. **Chance**: shuffling family labels or GenePT targets in train+val only should drive test scores to chance, which checks the pipeline for target leakage. **Protein upper bound**: ESM-2 (Lin et al., 2023) (650 M) on the protein sequence, mean-pooled over residues. **Supervised regulatory comparator**: Enformer (Avsec et al., 2021) on TSS windows (§2.2). It is trained on regulatory tracks, not family labels, so it is a reference point rather than a bound.

### 2.2 Evaluation axes

#### Pooling

Encoders read long genes in chunks that fit their context windows, so each gene must be reduced to one vector, and that reduction can change what a probe sees. Parameter-free rules then reduce the chunks (Table 1): standard reductions for contextual embeddings (Devlin et al., 2019; Reimers and Gurevych, 2019), two that also keep the first and last chunks, and, for TSS windows, TSS-Anchored, which keeps only the chunk nearest the TSS.

#### Substrate

Two substrates carry the same gene. **CDS**: the coding sequence encodes the protein, so it carries the family label directly. **TSS window**: 196,608 bp of Ensembl sequence centred on each gene’s strand-specific transcription start site, mostly noncoding, asking whether the label can also be read from the sequence around the gene’s start. On TSS windows we ran all four encoders, the pooling rules except Mean (Boundary-including), and a matched 4-mer baseline; Enformer is read both as its whole-window trunk mean and over its central *∼*2,048 bp.

#### Split A random split

assigns genes individually, stratified by family, and lets closely related genes (paralogs, copies of a gene created by duplication) fall on both sides of the partition: 56% of its test genes have a relative in training or validation, so a probe can score by recognising near-copies (Appendix C.4). We therefore evaluate on two homology-aware splits and keep the random split as a control (§3.5). The **homology split**, primary for CDS, translates each canonical CDS, clusters the proteins with MMseqs2 (Steinegger and Söding, 2017) at 40% identity and 80% bidirectional coverage, above Rost’s “twilight zone” (Li and Godzik, 2006; Rost, 1999), and assigns whole clusters, largest first, to one partition in 70/15/15 proportions stratified by family (Appendix Table A1); no test gene then shares a cluster with a training gene. The **disjoint split**, primary for TSS, additionally groups genes whose windows overlap on the genome, since windows of genes in different partitions can otherwise share sequence. Because MMseqs2 compares genes to a cluster representative rather than pairwise, a few close pairs still straddle the split; test genes with such a partner in training or validation (or, on the disjoint split, an overlapping window) are dropped from scoring, leaving 459 CDS and 465 TSS test genes. The homology split is repeated with four seeds for assigning clusters to partitions (Appendix Table C1; settings in Appendix D.1).

## 3 Results and Discussion

Differences are paired bootstrap estimates over groups of related test genes, with 95% intervals (Appendix Table C3; per-cell intervals in Appendix Tables B1 and B2; split-seed ranges in Appendix Table C1). Four comparisons are primary, one-sided and Holm-corrected together: the selected encoder against the nucleotide *k*-mer and against the amino-acid *k*-mer, ESM-2 against the selected encoder, all on the homology split (§3.1), and NT-v2 on CDS against the TSS window of the same genes on the disjoint split (§3.4). We fixed the four after seeing the submitted results on these test genes, so they are not pre-registered (Appendix E). The remaining sections quantify how much pooling, substrate and split move scores, with uncorrected *p*-values; the pooling ranges in §3.3 are point estimates, not tests.

### 3.1 CDS encoders reach nucleotide composition, not amino-acid composition

If frozen DNA encoders capture protein-level signal, they should beat nucleotide composition on CDS; if that signal goes no further than composition, they should not beat amino-acid composition. The selected encoder does not significantly beat nucleotide composition, and no encoder beats amino-acid composition. The validation-selected encoder, NT-v2, beats 6-mer composition by +0.054 macro-F1 [*−*0.011, +0.126], not significant after Holm correction (*p* = 0.108), and trails amino-acid 3-mer composition (*−*0.075 [*−*0.137, *−*0.001]). ESM-2, reading the protein directly, sits +0.261 [+0.200, +0.322] above NT-v2. GENA-LM scores highest on test (0.710) but also does not reach amino-acid composition (0.767). DNABERT-2 and HyenaDNA sit at the nucleotide floor (*−*0.011 and *−*0.006; Figure 2; Appendix Table B1).

**Figure 2:**
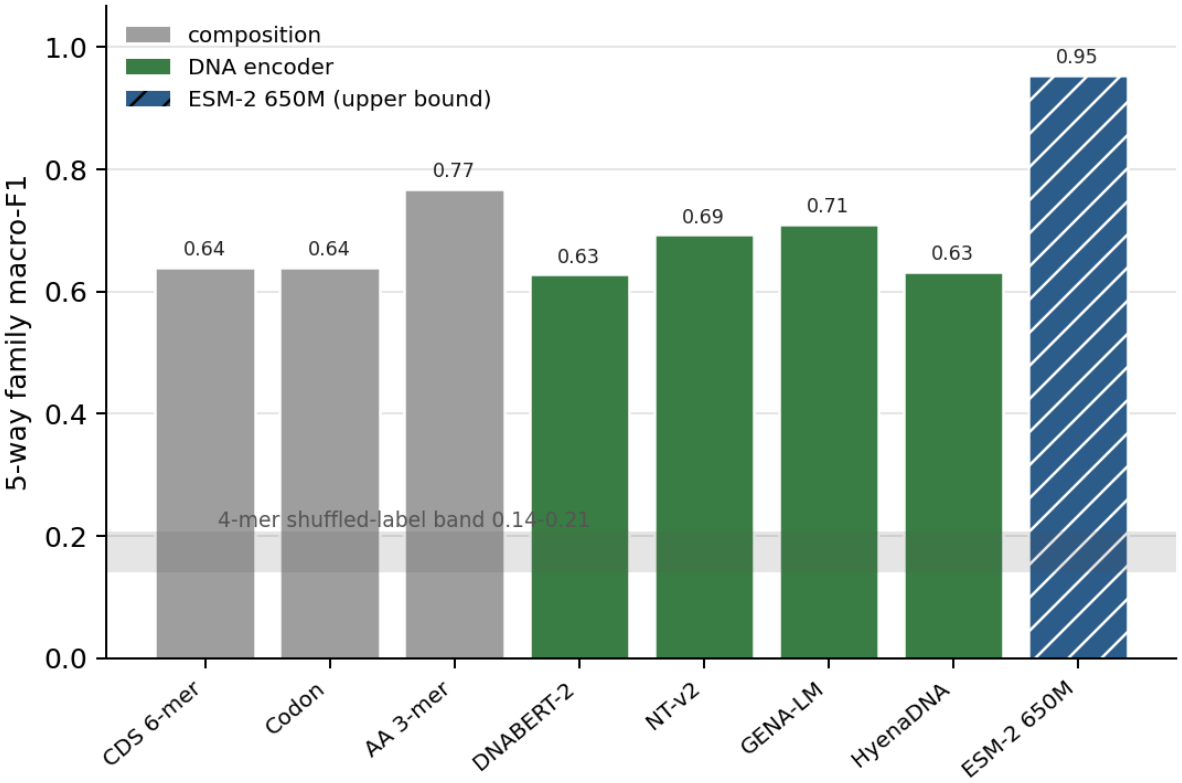
5-way family classification macro-F1 on CDS: composition baselines (grey), DNA encoders at their validation-selected pooling (green), and ESM-2 650M on the protein sequence (blue, hatched). The nucleotide and amino-acid k are chosen on validation.

We read this as NT-v2 pointing above nucleotide composition but not reliably (Holm-adjusted *p* = 0.108) and below translation, GENA-LM scoring similarly on test though it was not the validation pick, and DNABERT-2 and HyenaDNA recovering nothing beyond composition. Translated amino-acid counts use the reading frame the encoders are never given, and ESM-2 marks how much more a protein-native model reads.

### 3.2 Encoders recover little text-described function, and none beyond aminoacid composition

If the family signal reflected gene function beyond protein composition, it should transfer to a text-derived function target. It does not. The best encoder, DNABERT-2, reaches GenePT *R*^2^ 0.065: above nucleotide 6-mer composition (+0.026 [+0.018, +0.034]), but at or below amino-acid 3-mer composition (*−*0.015 [*−*0.031, +0.000]), and ESM-2 sits +0.096 above amino-acid composition (Figure 3; Appendix Table B2).

**Figure 3:**
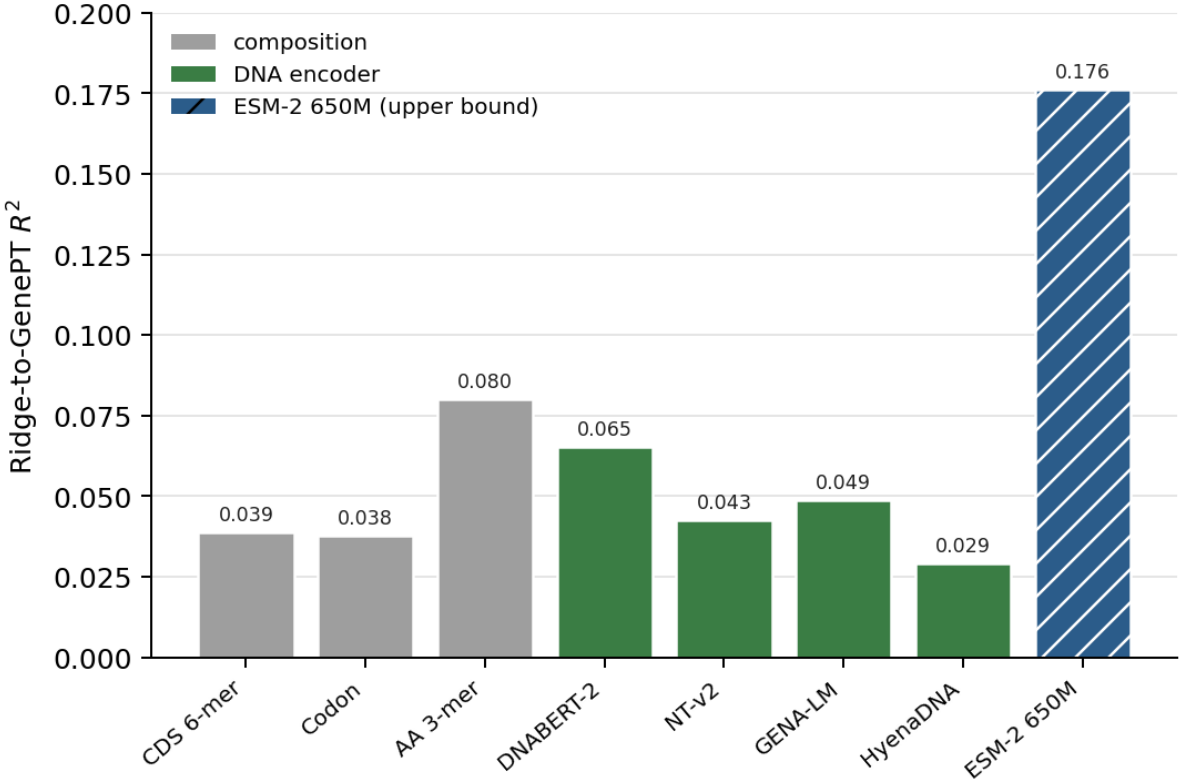
Ridge-to-GenePT R^2^ on CDS, averaged over the 1,536 GenePT dimensions, for the same features and colours as Figure 2. The nucleotide and amino-acid k are chosen on validation for this probe.

Part of the weak alignment is the target itself. GenePT vectors inherit the geometry of English gene summaries, which sequence does not determine, so all vectors point in similar directions. Cosine similarity therefore stays high even for shuffled targets and cannot separate the features; *R*^2^ can, which is why we report it, but even ESM-2 reaches only 0.176. Stronger alignment may require training it explicitly, as DNA–text models such as Omni-DNA do (Li et al., 2025, 2026).

### 3.3 Pooling can move an encoder from below the composition floor to level with it

If pooling were a neutral default, an encoder’s score would not depend on it. It does: NT-v2 scores macro-F1 0.504 with Max and 0.698 with Mean (Boundary-including), against 0.639 for the validation-selected 6-mer, so pooling alone moves it from well below the composition floor to level with it. Pooling moves a single encoder by 0.159 to 0.196 macro-F1, well above the 0.041 to 0.087 that separates the four encoders at any one rule they all share, and Max is the weakest rule for every encoder. Because the best rule is encoder-specific, each encoder is reported at its validation-selected rule (Figure 4; full matrix in Appendix Table B5). Pooling is therefore a first-order design choice: frozen-encoder comparisons need explicit pooling controls, not a single default.

**Figure 4:**
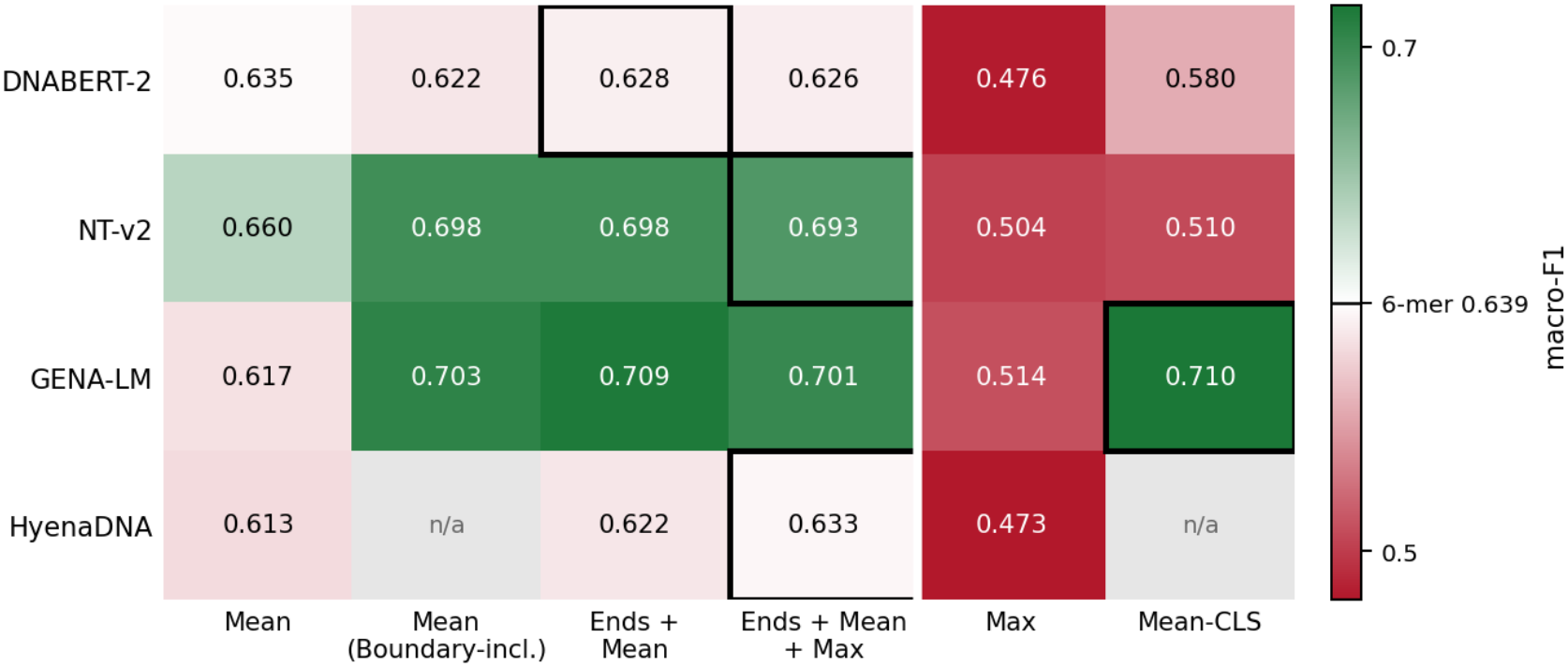
Macro-F1 across all encoder-pooling cells (CDS). Colour is centred on the validation-selected CDS 6-mer (white): green above, red below, darker further away. Boxes mark each encoder’s validation-selected rule; n/a: the rule reads a trained boundary token, which HyenaDNA lacks.

The rules that add the first and last chunks are mostly a regularisation effect: 81.9 to 99.0% of genes fit one window, where Ends + Mean is three copies of Mean. Against Mean repeated three times at matched regularisation, only GENA-LM keeps a lead (+0.086 [+0.017, +0.153]; Appendix Table C3).

### 3.4 The TSS window carries little of the family label

The TSS window is mostly promoter, intronic, and intergenic sequence (about 1% is the gene’s own CDS; Appendix C.3), so any signal should be regulatory rather than family. The disjoint split, which removes the window overlap that 46.2% of homology-split test genes have with a training or validation gene, lowers DNABERT-2 and NT-v2 but not the other models (Appendix Table C2). We read the window two ways, averaged over its whole length (§3.4.1) and at the TSS (§3.4.2), and Enformer the same two ways.

#### 3.4.1 Whole-window pooling

If the window carried the family label, whole-window averages should approach CDS scores on the same genes; if it carries only promoter composition, they should sit near the TSS 4-mer. The second holds. The encoder selected on TSS validation, DNABERT-2, reaches macro-F1 0.258, below the TSS 4-mer (0.281) but above chance (shuffled-label median 0.167); the best encoder on test, GENA-LM, reaches 0.340. Enformer, trained on regulatory tracks and averaged over its output bins, reaches 0.413. Every encoder falls far below its CDS score on the same test genes (CDS minus TSS +0.267 to +0.393; Appendix Table C3). The window has no reading frame, so amino-acid composition is undefined and the 4-mer is the only composition anchor; GenePT *R*^2^ is below zero for every TSS feature, so the window is reported on family macro-F1 only (Appendix Figure B1).

#### 3.4.2 TSS-Anchored pooling

If a family signal sits near the TSS and whole-window averaging dilutes it, the TSS-Anchored readout (Methods) should recover it. It does not, reliably. All four encoders gain a little from reading at the TSS (+0.012 to +0.050 macro-F1), but every paired interval includes zero; the largest gain, NT-v2’s, is +0.050 [*−*0.005, +0.106], and Enformer’s central readout matches its whole-window mean (*−*0.001). The composition of the anchored chunk alone does about as well: for three of four encoders the anchored score is within interval of the chunk’s own 4-mer or 6-mer baseline (diamonds in Figure 5; *−*0.049 to +0.015), and only GENA-LM beats it (+0.104 [+0.044, +0.161]). What the anchored readout recovers is therefore mostly the local composition around the TSS, not a diluted family signal, and every TSS result stays far below CDS (Appendix Tables B3 and C5).

**Figure 5:**
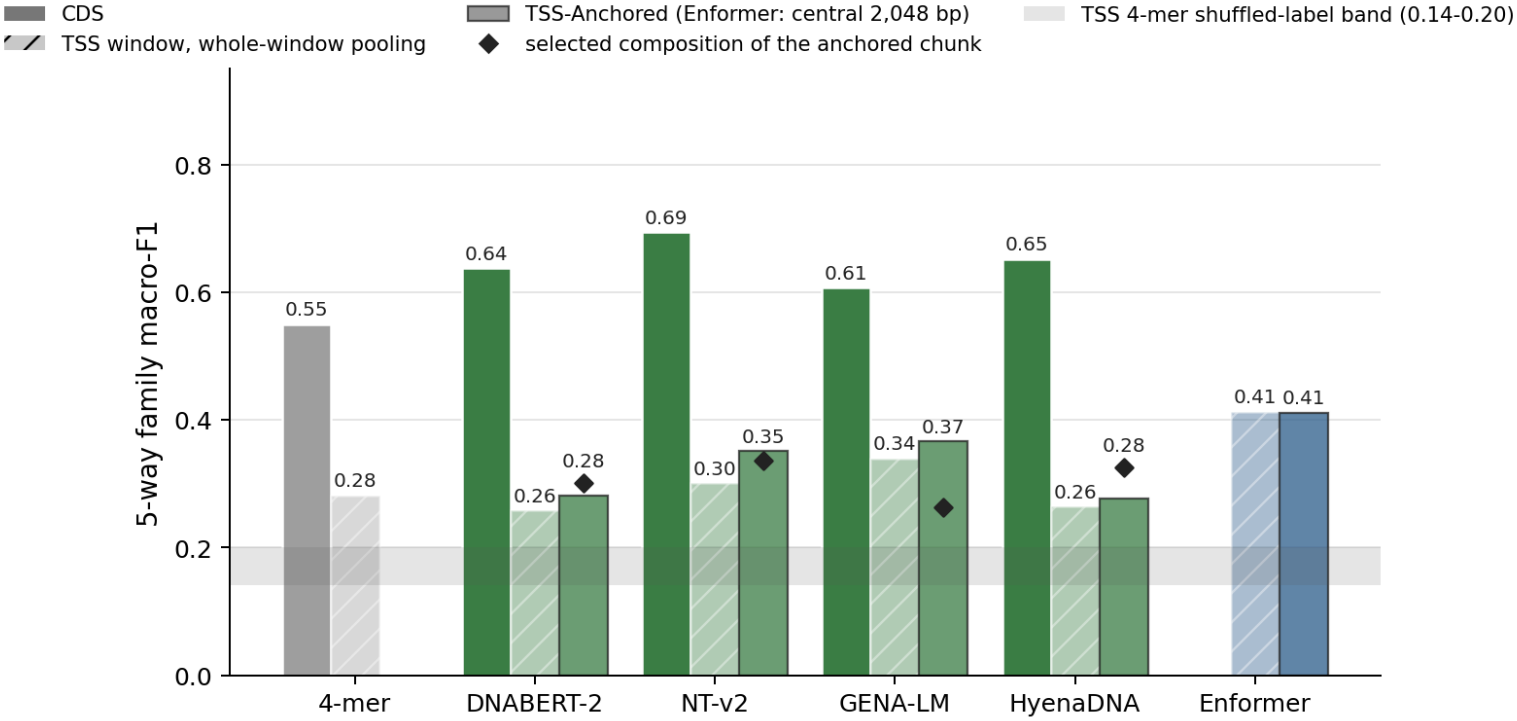
Family macro-F1 on CDS, the whole TSS window, and the TSS-Anchored chunk, all on the disjoint split. Enformer: whole-window mean and central 2,048 bp readout. Diamonds: the anchored chunk’s validation-selected composition baseline. GenePT R^2^ for the same cells is in Appendix Figure B1.

We interpret the TSS drop as a mismatch between substrate and target. The same frozen encoders recover the family label from CDS, so they can represent it; what the window lacks is its source, since only about 1% of it is the gene’s own CDS. Consistent with this, reading at the TSS adds no reliable signal over the whole window, and no readout, including the supervised Enformer, approaches CDS. Whether a learned attention pooling over frozen token embeddings, or fine-tuning, could find the coding exons inside the window, and whether a regulatory target can be read from it, we leave to future work.

### 3.5 Random splits infiate the TSS window more than CDS

If a random split leaks homologous genes into the test set, scores should be higher there than on each arm’s primary split. Re-running every comparator on a random split measures the gap. The central reading survives: amino-acid composition leads the best DNA encoder on both splits (0.908 vs. 0.801 random, 0.767 vs. 0.693 homology), and ESM-2 leads throughout. What changes is the level, and unevenly. Every model and baseline scores higher on the random split except GENA-LM on CDS, which ties (Appendix Table B4). Counting only the margin above chance (macro-F1 0.167 under shuffled labels), NT-v2 keeps 83% of its random-split margin on CDS under the homology split but only 42% on the TSS window under the disjoint split, so a random split infiates the regulatory arm far more than the coding arm (Figure 6; Appendix Table B4).

**Figure 6:**
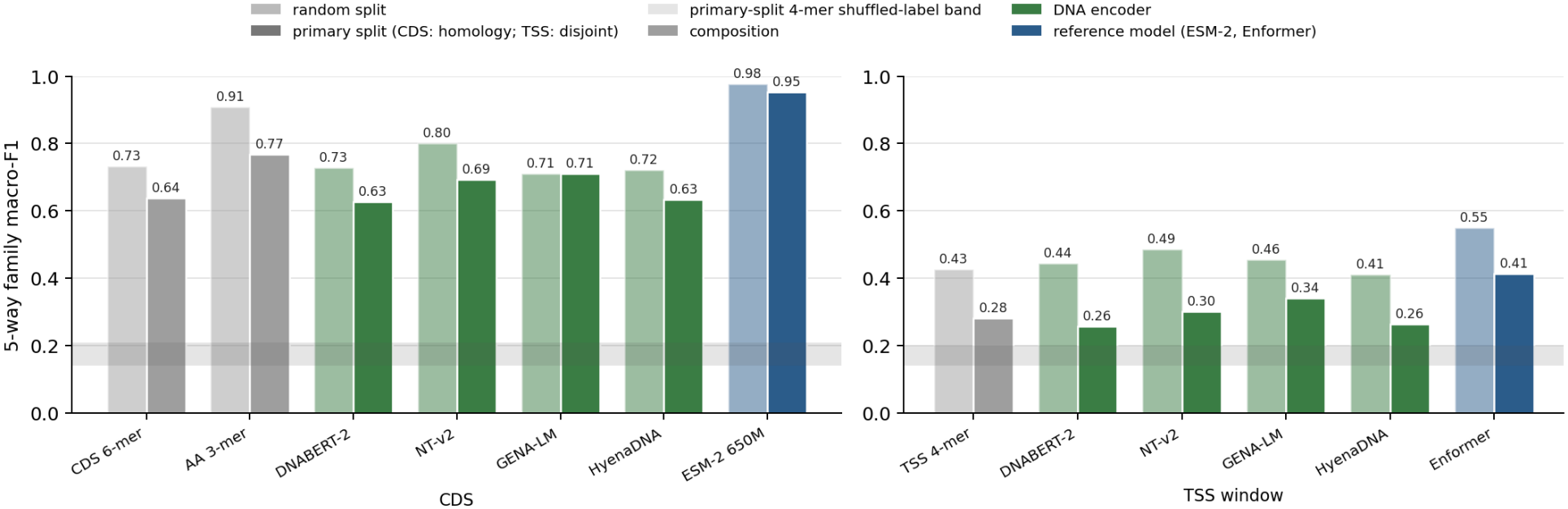
Family macro-F1 on the random-stratified split against each arm’s primary split: CDS on the homology-aware split (left) and the TSS window on the disjoint split (right), every model at its validation-selected pool on each split. Shaded: each primary split’s 4-mer shuffled-label band. GenePT R^2^ per split is in Appendix Table B4.

We read this as evidence that the homology split, not a composition baseline alone, is what separates memorised similarity from transferable signal, the coding-sequence counterpart of the leakage Rafi et al. (2025) report for DNA-level expression models.

## 4 Conclusion and Limitations

### 4.1 Conclusion

MINA fixes a task and varies how it is evaluated. On that task, read through linear probes, frozen DNA encoders recover coarse protein-family signal from coding sequence at about the level of nucleotide composition, do not reach translated amino-acid composition, and recover little of the text-described function in GenePT. How the task is evaluated changes the conclusions as much as which encoder is used: the pooling rule alone can move an encoder from well below the composition floor to level with it, a mostly noncoding substrate cuts the family signal far below CDS, and random splits infiate most scores, on CDS through homologous genes.

We draw four recommendations for evaluating frozen DNA embeddings: compare every score against nucleotide and amino-acid composition; treat pooling as a reported, validation-selected choice rather than a default; match the substrate to the target; and split by homology, and by genomic interval when inputs are windows. The next steps are a learned pooling over frozen token embeddings, regulatory targets for the TSS window, and families beyond the five used here.

### 4.2 Limitations

#### Scope

MINA is small by design: cheap to repeat across encoders, pooling rules, substrates and splits, at the cost of power and breadth. It covers 3,244 human genes in five broad HGNC families, 99.9% of GenePT-covered genes in those families; the remaining *∼*15,600 GenePT genes have no broad-family ontology, so a larger label set is a separate many-class study. The five families are a best case, large and domain-rich, and every comparison is within this one task with one chance floor, so the conclusions may not transfer to families with weaker signatures or to other class counts. The protocol is species-agnostic; GenePT and the HGNC families are not.

#### Labels and targets

Family labels come from broad HGNC group names, which add noise: 387 transcription-factor labels come only from zinc-finger groups outside the C2H2 class (a rule that also catches genuine factors such as GATA1–6), and 150 kinase labels are not protein kinases; masking either set leaves every primary test pointing the same way (Appendix Table C3). GenePT is the only text target and enters only the regression arm. It inherits literature bias, and 901 genes share a templated summary; masking them leaves the ordering unchanged. Macro-averaged *R*^2^ depends on the GenePT basis; pooled *R*^2^, which does not, keeps the encoder ordering, and retrieval@5 reorders the DNA encoders but keeps ESM-2 first (Appendix Table C4).

#### Readout

A linear probe on a frozen, pooled embedding is a lower bound on what the embedding contains: signal it finds is there, but signal it misses may still be recoverable by a nonlinear probe, learned pooling, or fine-tuning. The DNA encoders were pretrained on unlabelled genomes, ESM-2 on UniRef proteins, and Enformer on human and mouse regulatory tracks, so each may have seen our test sequences; none was trained on the family labels or GenePT targets.

#### Statistics

The test set is small: 459 CDS genes, only 22 of them immune receptors, so a paired macro-F1 interval spans about *±*0.07 and smaller differences are not resolved, and macro-F1 lets those 22 genes count as much as the 245 transcription factors. The intervals cover only the sampling of test genes, not the uncertainty from hyperparameter, split-seed, or pooling choices, which we report separately (split seeds in Appendix Table C1, pooling in Appendix Table B5, the numerical backend in Appendix D.1). TSS-Anchored pooling was designed after seeing the whole-window results, so it is exploratory.

## Acknowledgements

We thank Professor Cassandra Burdziak for helpful feedback and guidance during the development of this project. This work received no external funding. The authors declare no competing interests.

