## Appendix for "MINA: Do frozen DNA encoders beat sequence composition? A homology-aware linear-probe benchmark"

### A Dataset, Baselines and Encoders

Appendix A defines the dataset, baselines and encoders; Appendix B gives the full values behind every main-text figure and the complete pooling and Ridge grids; Appendix C reports confidence intervals and robustness checks; Appendix D covers code and settings; Appendix E lists what changed since submission. CDS tables use the homology-aware split; TSS tables and every CDS–TSS comparison use the disjoint split (both 70/15/15). Random-split tables say so. Every table uses the main text's validation-tuned probe protocol. A dagger (†) marks a digit that is not stable across numerical backends of the solver (Appendix D.1); no primary test is affected.

#### A.1 Dataset Composition and Baseline Definitions

Table A1: Curation. Each family's include patterns are matched, case-insensitively, against the HGNC `gene_group` field of protein-coding genes with an Ensembl ID; exclude patterns (in parentheses) remove a gene from that family. Match: genes matching; GenePT: also in GenePT; Kept: after first-family-wins assignment in the order shown (17 genes match two families); Train/Val/Test: the homology-aware split. Three olfactory receptors have no retrieved CDS, so GPCR enters the split with 591 and the dataset holds 3,244.

| Family | Include (exclude) patterns | Match | GenePT | Kept | Train | Val | Test |
|---|---|---|---|---|---|---|---|
| Kinase | `\bkinase` (kinase inhibitor, kinase regulator, kinase substrate, pseudokinase) | 564 | 558 | 558 | 391 | 83 | 84 |
| TF | zinc finger, homeobox, basic helix-loop-helix, `\bbZIP\b`, forkhead box, high mobility group, nuclear receptor, T-box, SOX transcription, ETS transcription, transcription factor (transcription factor binding, cofactor) | 1,757 | 1,745 | 1,743 | 1,220 | 262 | 261 |
| Ion channel | `\bchannel` (channel regulat, channel auxiliary, channel interacting) | 201 | 200 | 198 | 138 | 30 | 30 |
| GPCR | G protein-coupled receptor, adrenoceptor, 5-hydroxytryptamine receptor, dopamine receptor, muscarinic, opioid receptor, chemokine receptor, olfactory receptor | 617 | 601 | 594 | 414 | 88 | 89 |
| Immune | toll like receptor, `interleukin .* receptor`, `\bFc receptor`, NOD-like receptor, NLR family, killer cell immunoglobulin, T cell receptor, B cell receptor, C-type lectin domain, immunoglobulin like receptor (binding) | 164 | 160 | 154 | 108 | 23 | 23 |

Table A2: Parameter-free composition baselines, each an $L_1$-normalised frequency vector run through the same probe recipe. Controls append raw log-length (and GC) terms and are never candidates for the headline composition pick: the length controls feed the composition-with-length sensitivity test (Table C3), and the chunk baselines are the composition of the same anchored chunk an encoder reads (Table C5).

| Baseline | Feature | Dim |
|---|---|---|
| CDS 4-mer | Overlapping 4-nucleotide windows, frame-agnostic | 256 |
| CDS 6-mer | Overlapping 6-nucleotide windows | 4,096 |
| Codon | In-frame codons (frame 0, non-overlapping) | 64 |
| AA 1-mer | Amino-acid composition of translated CDS | 20 |
| AA 2-mer | Overlapping dipeptide frequencies | 400 |
| AA 3-mer | Overlapping tripeptide frequencies | 8,000 |
| *Controls* | | |
| · + length | A nucleotide or AA $k$-mer above, plus $\log(1 + \text{CDS length})$ | +1 |
| Chunk 4-mer + GC | Anchored chunk: 4-mer, GC, two log-lengths | 259 |
| Chunk 6-mer | Anchored chunk: 6-mer | 4,096 |

##### A.2 Encoder Architectures

**DNABERT-2** (Zhou *et al.*, 2024) is a 117 M-parameter BERT-style model (`zhihan1996/DNABERT-2-117M`) with byte-pair encoding, 768-d hidden states, and a 512-token context window. **NT-v2** (Dalla-Torre *et al.*, 2025) (`InstaDeepAI/nucleotide-transformer-v2-100m-multi-species`) is an ESM-style masked-language model with fixed non-overlapping 6-mer tokens, 512-d hidden states, and a 2,048-token context window. **GENA-LM base** (Fishman *et al.*, 2025) (`AIRI-Institute/gena-lm-bert-base-t2t`) is a 110 M-parameter BERT-12L masked-language model with 768-d hidden states, 12 attention heads, a 32,000-token BPE vocabulary, and a 512-token context window corresponding to roughly 4.5 kb after BPE compression; the `t2t` checkpoint was trained on the human T2T assembly with 1000 Genomes SNP augmentation. **HyenaDNA large** (Nguyen *et al.*, 2023) (`LongSafari/hyenadna-large-1m-seqlen-hf`) is an attention-free causal next-nucleotide model based on Hyena sequence operators, with single-nucleotide tokens, 8 layers, 256-d hidden states, approximately 6.6 M trainable parameters, and a context window up to 1M nucleotides; HyenaDNA was pretrained on the human reference genome.

### B Full Values

#### B.1 Full Values for Main-Text Figures

Each table below gives the exact values plotted in the corresponding main-text figure.

Table B1: Full values for Figure 2 (5-way family classification), each with its 95% cluster-bootstrap CI (1,000 resamples); the shuffled-label row carries its null band. Macro-F1 and $\kappa$ order neighbouring encoders differently but agree that no DNA encoder reaches amino-acid composition. Bold: the best value in each score column outside ESM-2 (the upper bound) and the controls.

| Source | Pool | F1 [95% CI] | ΔF1 | $\kappa$ [95% CI] | Acc. |
|---|---|---|---|---|---|
| Shuffled labels (median of 200) | — | 0.167 [0.141, 0.208] | -0.472 | -0.015 | 0.475 |
| CDS 6-mer | — | 0.639 [0.572, 0.697] | +0.000 | 0.593 [0.521, 0.653] | 0.741 |
| Codon | — | 0.640 [0.569, 0.704] | +0.001 | 0.617 [0.550, 0.682] | 0.767 |
| AA 1-mer | — | 0.697 [0.637, 0.761] | +0.058 | 0.679 [0.616, 0.739] | 0.800 |
| AA 2-mer | — | 0.760 [0.698, 0.816] | +0.122 | 0.729 [0.668, 0.783] | 0.832 |
| AA 3-mer | — | **0.767 [0.703, 0.821]** | +0.129 | **0.757 [0.698, 0.808]** | **0.847** |
| DNABERT-2 | Ends + Mean | 0.628 [0.557, 0.690] | -0.011 | 0.598 [0.536, 0.666] | 0.749 |
| NT-v2 | Ends + Mean + Max | 0.693 [0.629, 0.754] | +0.054 | 0.653 [0.594, 0.710] | 0.780 |
| GENA-LM | Mean-CLS | 0.710 [0.647, 0.765] | +0.071 | 0.651 [0.590, 0.710] | 0.782 |
| HyenaDNA | Ends + Mean + Max | 0.633 [0.571, 0.686] | -0.006 | 0.573 [0.503, 0.631] | 0.730 |
| ESM-2 650M | — | 0.954 [0.923, 0.977] | +0.315 | 0.939 [0.907, 0.966] | 0.961 |

Table B2: Full values for Figure 3 (Ridge-to-GenePT regression), each with its 95% cluster-bootstrap CI. Bold: the best value in each score column outside ESM-2 (the upper bound) and the controls.

| Source | Pool | $R^2$ [95% CI] | Δ | Cos. |
|---|---|---|---|---|
| CDS 6-mer | — | 0.039 [0.014, 0.057] | +0.000 | 0.916 |
| Codon | — | 0.038 [0.018, 0.048] | -0.001 | 0.916 |
| AA 1-mer | — | 0.022 [0.004, 0.033] | -0.017 | 0.915 |
| AA 2-mer | — | 0.059 [0.040, 0.073] | +0.021 | 0.918 |
| AA 3-mer | — | **0.080 [0.058, 0.094]** | +0.042 | **0.920** |
| DNABERT-2 | Mean (Boundary-including) | 0.065 [0.038, 0.089] | +0.026 | 0.919 |
| NT-v2 | Mean (Boundary-including) | 0.043 [0.018, 0.057] | +0.004 | 0.917 |
| GENA-LM | Ends + Mean | 0.049 [0.024, 0.067] | +0.010 | 0.917 |
| HyenaDNA | Ends + Mean | 0.029 [0.006, 0.044] | -0.010 | 0.915 |
| ESM-2 650M | — | 0.176 [0.152, 0.191] | +0.137 | 0.929 |

Table B3: Full values for the CDS and whole-window bars of Figure 5: per-encoder classification (macro-F1, with $\kappa$) and Ridge-to-GenePT ($R^2$), CDS versus TSS window under whole-window pooling. Every row is on the disjoint split, so CDS and TSS score the same test genes; Δ is against the 4-mer of the same substrate, the only composition baseline defined on the window (Table B1 uses the validation-selected $k$ on the homology split). On CDS the stronger baselines are defined too (6-mer 0.627, amino-acid 3-mer 0.762), and every encoder's macro-F1 stays below amino-acid composition, as on the homology split. Bold: the best DNA encoder's F1 and ΔF1 in each block, and its CDS $R^2$ and $\Delta R^2$; TSS $R^2$ is below zero for every encoder and is not bolded.

| Source | F1 | ΔF1 | $\kappa$ | $R^2$ | $\Delta R^2$ |
|---|---|---|---|---|---|
| **Coding sequence (CDS)** | | | | | |
| 4-mer | 0.550 | +0.000 | 0.509 | 0.014 | +0.000 |
| DNABERT-2 | 0.638 | +0.088 | 0.607 | **0.048** | **+0.034** |
| NT-v2 | **0.694** | **+0.144** | 0.644 | 0.025 | +0.011 |
| GENA-LM | 0.607$^\dagger$ | +0.057$^\dagger$ | 0.537$^\dagger$ | 0.027 | +0.013 |
| HyenaDNA | 0.652 | +0.102 | 0.587 | 0.015 | +0.001 |
| **TSS-centred window (196,608 bp)** | | | | | |
| 4-mer | 0.281 | +0.000 | 0.123 | -0.034 | +0.000 |
| DNABERT-2 | 0.258 | -0.024 | 0.104 | -0.011 | +0.023 |
| NT-v2 | 0.300 | +0.019 | 0.190 | -0.013 | +0.021 |
| GENA-LM | **0.340** | **+0.058** | 0.192 | -0.020 | +0.014 |
| HyenaDNA | 0.264 | -0.017 | 0.113 | -0.037 | -0.003 |
| Enformer (whole window) | 0.413 | +0.131 | 0.268 | -0.006 | +0.028 |

Table B4: Full values for Figure 6: random-stratified versus each arm's primary split (CDS homology-aware, TSS disjoint), by sequence context, with leakage deltas.

| Source | F1 (rand) | F1 (prim.) | ΔF1 | $R^2$ (rand) | $R^2$ (prim.) | $\Delta R^2$ |
|---|---|---|---|---|---|---|
| **Coding sequence (CDS)** | | | | | | |
| CDS 6-mer | 0.733 | 0.639 | -0.094 | 0.190 | 0.039 | -0.151 |
| Codon | 0.693 | 0.640 | -0.054 | 0.173 | 0.038 | -0.135 |
| AA 2-mer | 0.854 | 0.760 | -0.094 | 0.204 | 0.059 | -0.144 |
| AA 3-mer | 0.908 | 0.767 | -0.141 | 0.278 | 0.080 | -0.198 |
| NT-v2 | 0.801 | 0.693 | -0.108 | 0.192 | 0.043 | -0.149 |
| DNABERT-2 | 0.728 | 0.628 | -0.101 | 0.210 | 0.065 | -0.145 |
| GENA-LM | 0.710 | 0.710 | -0.000 | 0.191 | 0.049 | -0.142 |
| HyenaDNA | 0.721 | 0.633 | -0.089 | 0.181 | 0.029 | -0.152 |
| ESM-2 650M | 0.977 | 0.954 | -0.023 | 0.350 | 0.176 | -0.174 |
| **TSS window (196,608 bp; primary = genomic-interval-disjoint split)** | | | | | | |
| TSS 4-mer | 0.427 | 0.281 | -0.145 | 0.086 | -0.034 | -0.120 |
| DNABERT-2 | 0.445 | 0.258 | -0.187 | 0.129 | -0.011 | -0.140 |
| NT-v2 | 0.487 | 0.300 | -0.187 | 0.124 | -0.013 | -0.137 |
| GENA-LM | 0.456 | 0.340 | -0.116 | 0.113 | -0.020 | -0.133 |
| HyenaDNA | 0.410 | 0.264 | -0.146 | 0.091 | -0.037 | -0.128 |
| Enformer (whole window) | 0.550 | 0.413 | -0.137 | 0.134 | -0.006 | -0.140 |

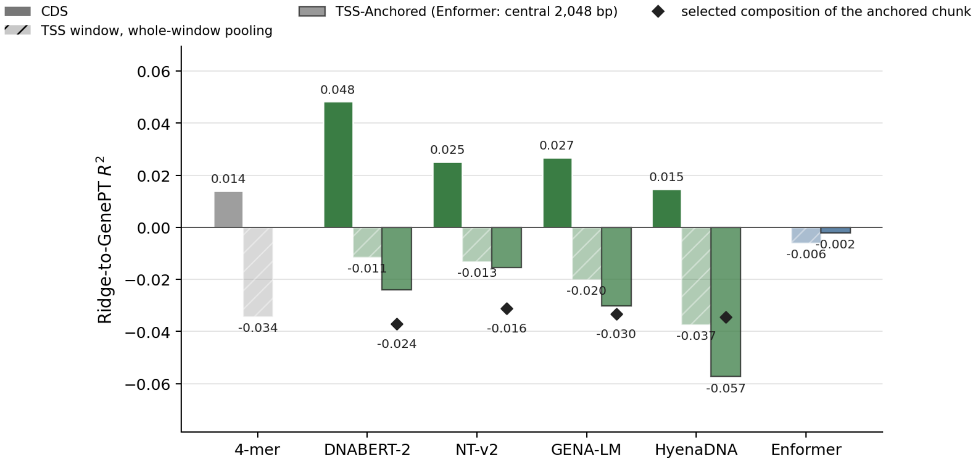


*Figure B1: Ridge-to-GenePT $R^2$ for the cells of Figure 5: CDS, the whole TSS window, and the TSS-Anchored chunk, on the disjoint split. Every TSS feature scores below zero, so the probe predicts the text embedding worse than the training mean does; the main text therefore reports the TSS window on family macro-F1 only. CDS and whole-window values are in Table B3; TSS-Anchored values are printed on the bars.*

#### B.2 Full pooling and Ridge matrices: primary vs. random split

Each readout is shown as one full-width matrix with the primary split (homology-aware for CDS, disjoint for TSS) beside its random-stratified mirror, so any encoder-pooling cell can be read across the two splits to gauge the homology-leakage inflation. Every cell's macro-F1 is higher on the random split except GENA-LM's CDS Mean-CLS, Ends + Mean and Ends + Mean + Max cells, and translated amino-acid composition matches or beats the best DNA encoder on both. The full classification matrix is Table B5 and the full Ridge matrix is Table B6, given on the following pages.

Table B5: Every cell for 5-way family classification: the primary split (left; CDS on the homology-aware split, TSS on the genomic-interval-disjoint split) versus the random-stratified split (right). Each split selects $C$ on its own validation set. Mean and Mean (Boundary-including) differ only by the boundary tokens: for NT-v2 and GENA-LM they score alike wherever validation picks the same $C$, and on the primary CDS split validation picked a far larger $C$ for Mean on a nearly flat curve, which costs it on test.

| | | **Primary split** | | | **Random-stratified split** | | |
|---|---|---|---|---|---|---|---|
| Source | Pooling | Macro-F1 | $\kappa$ | Accuracy | Macro-F1 | $\kappa$ | Accuracy |
| **Coding sequence (CDS; primary = homology-aware split)** | | | | | | | |
| CDS 4-mer | — | 0.6482 | 0.5905 | 0.7495 | 0.6852 | 0.7018 | 0.8090 |
| CDS 6-mer | — | 0.6386 | 0.5927 | 0.7407 | 0.7327 | 0.7314 | 0.8275 |
| Codon | — | 0.6397 | 0.6165 | 0.7669 | 0.6932 | 0.7154 | 0.8214 |
| AA 1-mer | — | 0.6970 | 0.6792 | 0.7996 | 0.7064 | 0.7214 | 0.8255 |
| AA 2-mer | — | 0.7603 | 0.7289 | 0.8322 | 0.8540 | 0.8261 | 0.8891 |
| AA 3-mer | — | 0.7674 | 0.7570 | 0.8475 | 0.9081 | 0.9040 | 0.9384 |
| DNABERT-2 | Mean | 0.6351 | 0.6073 | 0.7538 | 0.7150 | 0.6976 | 0.8090 |
| DNABERT-2 | Mean (Boundary-including) | 0.6217 | 0.5991 | 0.7495 | 0.7205 | 0.7043 | 0.8131 |
| DNABERT-2 | Max | 0.4765 | 0.3899 | 0.6013 | 0.5875 | 0.5654 | 0.7228 |
| DNABERT-2 | Mean-CLS | 0.5798† | 0.5150† | 0.6797† | 0.6487 | 0.6693 | 0.7967 |
| DNABERT-2 | Ends + Mean | 0.6276 | 0.5983 | 0.7495 | 0.7361 | 0.7250 | 0.8255 |
| DNABERT-2 | Ends + Mean + Max | 0.6264 | 0.6005 | 0.7516 | 0.7284 | 0.7192 | 0.8234 |
| NT-v2 | Mean | 0.6604† | 0.6059† | 0.7495† | 0.8007 | 0.7947 | 0.8706 |
| NT-v2 | Mean (Boundary-including) | 0.6983 | 0.6577 | 0.7843 | 0.8007 | 0.7947 | 0.8706 |
| NT-v2 | Max | 0.5036 | 0.4987 | 0.7015 | 0.6038 | 0.6162 | 0.7659 |
| NT-v2 | Mean-CLS | 0.5105 | 0.4806 | 0.6754 | 0.6041 | 0.5782 | 0.7331 |
| NT-v2 | Ends + Mean | 0.6982 | 0.6633 | 0.7865 | 0.8108 | 0.8087 | 0.8789 |
| NT-v2 | Ends + Mean + Max | 0.6928 | 0.6525 | 0.7800 | 0.8028 | 0.7871 | 0.8645 |
| GENA-LM | Mean | 0.6172† | 0.5652† | 0.7168† | 0.7103 | 0.6954 | 0.8090 |
| GENA-LM | Mean (Boundary-including) | 0.7033 | 0.6450 | 0.7778 | 0.7103 | 0.6954 | 0.8090 |
| GENA-LM | Max | 0.5143 | 0.4120 | 0.6187 | 0.6002 | 0.5898 | 0.7474 |
| GENA-LM | Mean-CLS | 0.7100 | 0.6506 | 0.7821 | 0.6726 | 0.6864 | 0.8049 |
| GENA-LM | Ends + Mean | 0.7085 | 0.6597 | 0.7887 | 0.6627 | 0.6906 | 0.8111 |
| GENA-LM | Ends + Mean + Max | 0.7010 | 0.6584 | 0.7865 | 0.6984 | 0.7172 | 0.8234 |
| HyenaDNA | Mean | 0.6131 | 0.5595 | 0.7233 | 0.7025 | 0.6843 | 0.8049 |
| HyenaDNA | Max | 0.4730 | 0.3794 | 0.6122 | 0.5639 | 0.5304 | 0.7084 |
| HyenaDNA | Ends + Mean | 0.6218 | 0.5629 | 0.7233 | 0.7241 | 0.6894 | 0.8070 |
| HyenaDNA | Ends + Mean + Max | 0.6326 | 0.5727 | 0.7298 | 0.7214 | 0.6840 | 0.8029 |
| ESM-2 650M | — | 0.9536 | 0.9392 | 0.9608 | 0.9768 | 0.9745 | 0.9836 |
| **TSS-centred window (196,608 bp; primary = genomic-interval-disjoint split)** | | | | | | | |
| TSS 4-mer | — | 0.2814 | 0.1227 | 0.4817 | 0.4268 | 0.3503 | 0.6119 |
| DNABERT-2 | Mean | 0.2986 | 0.1332 | 0.5075 | 0.4446 | 0.3634 | 0.6099 |
| DNABERT-2 | Max | 0.2179 | 0.0962 | 0.5054 | 0.3940 | 0.2908 | 0.5688 |
| DNABERT-2 | Mean-CLS | 0.2861 | 0.1131 | 0.4731 | 0.3963 | 0.2769 | 0.5195 |
| DNABERT-2 | Ends + Mean | 0.2552 | 0.1452 | 0.4989 | 0.4223† | 0.3225† | 0.5914† |
| DNABERT-2 | Ends + Mean + Max | 0.2577 | 0.1043 | 0.4688 | 0.4137† | 0.3305† | 0.5749† |
| NT-v2 | Mean | 0.3004 | 0.1900 | 0.5419 | 0.4870 | 0.4217 | 0.6468 |
| NT-v2 | Max | 0.2515 | 0.1062 | 0.4774 | 0.3842 | 0.3234 | 0.5934 |
| NT-v2 | Mean-CLS | 0.2372 | 0.0763 | 0.4301 | 0.3107 | 0.2371 | 0.5791 |
| NT-v2 | Ends + Mean | 0.2941 | 0.1486 | 0.4731 | 0.3904 | 0.2873 | 0.5647 |
| NT-v2 | Ends + Mean + Max | 0.2873 | 0.1674 | 0.4839 | 0.3686 | 0.2789 | 0.5298 |
| GENA-LM | Mean | 0.3397 | 0.1916 | 0.5290 | 0.4557 | 0.3654 | 0.6160 |
| GENA-LM | Max | 0.2506 | 0.0690 | 0.4129 | 0.4168 | 0.3266 | 0.5975 |
| GENA-LM | Mean-CLS | 0.3398† | 0.1523† | 0.4323 | 0.4394† | 0.3255† | 0.5626† |
| GENA-LM | Ends + Mean | 0.3085 | 0.1865 | 0.5204 | 0.4145 | 0.3114 | 0.5893 |
| GENA-LM | Ends + Mean + Max | 0.2479† | 0.0990† | 0.4301† | 0.4387 | 0.3424 | 0.5975 |
| HyenaDNA | Mean | 0.2640 | 0.1132 | 0.4839 | 0.4150 | 0.3323 | 0.6119 |
| HyenaDNA | Max | 0.1947 | 0.0718 | 0.5097 | 0.3119 | 0.2310 | 0.5400 |
| HyenaDNA | Ends + Mean | 0.2638 | 0.1167 | 0.4645 | 0.3663 | 0.2407 | 0.5154 |
| HyenaDNA | Ends + Mean + Max | 0.2892 | 0.1730 | 0.5376 | 0.4103 | 0.2804 | 0.5606 |
| Enformer (whole window) | — | 0.4127 | 0.2677 | 0.5613 | 0.5498 | 0.4365 | 0.6427 |

Table B6: Every Ridge-to-GenePT cell: the primary split (left; CDS on the homology-aware split, TSS on the genomic-interval-disjoint split) versus the random-stratified split (right). Each split selects $\alpha$ on its own validation set.

| | | **Primary split** | | | **Random-stratified split** | | |
|---|---|---|---|---|---|---|---|
| Source | Pooling | $R^2$ macro | Mean cosine | $\alpha$ | $R^2$ macro | Mean cosine | $\alpha$ |
| **Coding sequence (CDS; primary = homology-aware split)** | | | | | | | |
| CDS 4-mer | — | 0.0320 | 0.9157 | 10000 | 0.1711 | 0.9304 | 1000 |
| CDS 6-mer | — | 0.0387 | 0.9165 | 10000 | 0.1900 | 0.9322 | 10000 |
| Codon | — | 0.0377 | 0.9165 | 1000 | 0.1732 | 0.9305 | 1000 |
| AA 1-mer | — | 0.0220 | 0.9147 | 10000 | 0.1662 | 0.9298 | 100 |
| AA 2-mer | — | 0.0595 | 0.9183 | 10000 | 0.2036 | 0.9333 | 10000 |
| AA 3-mer | — | 0.0802 | 0.9203 | 10000 | 0.2777 | 0.9399 | 10000 |
| DNABERT-2 | Mean | 0.0651 | 0.9188 | 10000 | 0.2062 | 0.9336 | 10000 |
| DNABERT-2 | Mean (Boundary-including) | 0.0652 | 0.9188 | 10000 | 0.2061 | 0.9336 | 10000 |
| DNABERT-2 | Max | 0.0398 | 0.9163 | 10000 | 0.1700 | 0.9302 | 10000 |
| DNABERT-2 | Mean-CLS | 0.0499 | 0.9174 | 10000 | 0.1880 | 0.9319 | 10000 |
| DNABERT-2 | Ends + Mean | 0.0640 | 0.9188 | 10000 | 0.2101 | 0.9340 | 10000 |
| DNABERT-2 | Ends + Mean + Max | 0.0599 | 0.9183 | 100000 | 0.2095 | 0.9339 | 10000 |
| NT-v2 | Mean | 0.0426 | 0.9169 | 1000 | 0.1916 | 0.9322 | 1000 |
| NT-v2 | Mean (Boundary-including) | 0.0426 | 0.9169 | 1000 | 0.1915 | 0.9322 | 1000 |
| NT-v2 | Max | 0.0205 | 0.9147 | 10000 | 0.1382 | 0.9271 | 10000 |
| NT-v2 | Mean-CLS | 0.0010 | 0.9126 | 10000 | 0.1168 | 0.9251 | 1000 |
| NT-v2 | Ends + Mean | 0.0416 | 0.9168 | 10000 | 0.1918 | 0.9323 | 1000 |
| NT-v2 | Ends + Mean + Max | 0.0426 | 0.9169 | 10000 | 0.1871 | 0.9318 | 10000 |
| GENA-LM | Mean | 0.0423 | 0.9167 | 10000 | 0.1824 | 0.9314 | 10000 |
| GENA-LM | Mean (Boundary-including) | 0.0423 | 0.9167 | 10000 | 0.1824 | 0.9314 | 10000 |
| GENA-LM | Max | 0.0223 | 0.9148 | 10000 | 0.1551 | 0.9288 | 10000 |
| GENA-LM | Mean-CLS | 0.0370 | 0.9162 | 10000 | 0.1885 | 0.9320 | 1000 |
| GENA-LM | Ends + Mean | 0.0488 | 0.9174 | 10000 | 0.1899 | 0.9321 | 10000 |
| GENA-LM | Ends + Mean + Max | 0.0493 | 0.9175 | 10000 | 0.1906 | 0.9322 | 10000 |
| HyenaDNA | Mean | 0.0263 | 0.9152 | 1000 | 0.1811 | 0.9312 | 100 |
| HyenaDNA | Max | -0.0015 | 0.9124 | 10000 | 0.1294 | 0.9263 | 1000 |
| HyenaDNA | Ends + Mean | 0.0290 | 0.9155 | 1000 | 0.1763 | 0.9307 | 1000 |
| HyenaDNA | Ends + Mean + Max | 0.0291 | 0.9155 | 1000 | 0.1780 | 0.9309 | 1000 |
| ESM-2 650M | — | 0.1760 | 0.9293 | 1000 | 0.3503 | 0.9464 | 1000 |
| **TSS-centred window (196,608 bp; primary = genomic-interval-disjoint split)** | | | | | | | |
| TSS 4-mer | — | -0.0343 | 0.9091 | 1000 | 0.0856 | 0.9218 | 1000 |
| DNABERT-2 | Mean | -0.0114 | 0.9111 | 1000 | 0.1288 | 0.9261 | 1000 |
| DNABERT-2 | Max | -0.0413 | 0.9083 | 1000 | 0.0875 | 0.9221 | 1000 |
| DNABERT-2 | Mean-CLS | -0.0236 | 0.9099 | 10000 | 0.0956 | 0.9230 | 1000 |
| DNABERT-2 | Ends + Mean | -0.0253 | 0.9098 | 10000 | 0.0934 | 0.9227 | 10000 |
| DNABERT-2 | Ends + Mean + Max | -0.0203 | 0.9103 | 10000 | 0.1067 | 0.9240 | 10000 |
| NT-v2 | Mean | -0.0131 | 0.9109 | 1000 | 0.1238 | 0.9257 | 100 |
| NT-v2 | Max | -0.0351 | 0.9089 | 1000 | 0.0837 | 0.9217 | 1000 |
| NT-v2 | Mean-CLS | -0.0347 | 0.9089 | 10000 | 0.0466 | 0.9181 | 1000 |
| NT-v2 | Ends + Mean | -0.0265 | 0.9097 | 10000 | 0.0771 | 0.9214 | 1000 |
| NT-v2 | Ends + Mean + Max | -0.0236 | 0.9100 | 10000 | 0.0847 | 0.9217 | 10000 |
| GENA-LM | Mean | -0.0171 | 0.9106 | 1000 | 0.1125 | 0.9245 | 1000 |
| GENA-LM | Max | -0.0229 | 0.9100 | 10000 | 0.0888 | 0.9221 | 10000 |
| GENA-LM | Mean-CLS | -0.0201 | 0.9103 | 1000 | 0.1069 | 0.9239 | 1000 |
| GENA-LM | Ends + Mean | -0.0313 | 0.9093 | 10000 | 0.0809 | 0.9215 | 10000 |
| GENA-LM | Ends + Mean + Max | -0.0309 | 0.9094 | 10000 | 0.0847 | 0.9219 | 10000 |
| HyenaDNA | Mean | -0.0373 | 0.9087 | 100 | 0.0905 | 0.9224 | 10 |
| HyenaDNA | Max | -0.0330 | 0.9091 | 10000 | 0.0700 | 0.9203 | 1000 |
| HyenaDNA | Ends + Mean | -0.0352 | 0.9089 | 10000 | 0.0591 | 0.9193 | 1000 |
| HyenaDNA | Ends + Mean + Max | -0.0334 | 0.9091 | 10000 | 0.0567 | 0.9191 | 1000 |
| Enformer (whole window) | — | -0.0060 | 0.9118 | 100000 | 0.1340 | 0.9267 | 10000 |

### C Confidence Intervals and Robustness Checks

#### C.1 Seed sensitivity and headline intervals

Table C1: Headline cells across four homology cluster-to-split seeds (42, 1, 7, 123), min–max. Each seed re-selects encoder, pooling and amino-acid $k$ on validation; the TSS row keeps DNABERT-2 and uses the seed's disjoint split. Seeds re-assign the same clusters largest-first, so two seed splits share 34–38% of CDS and 39–50% of TSS test genes (independent splits would share about 15%), so the ranges understate split-to-split variation.

| Cell | Macro-F1 (4 splits) | GenePT $R^2$ (4 splits) |
|---|---:|---:|
| ESM-2 650M | 0.926–0.955 | 0.176–0.191 |
| Best DNA encoder | 0.619$^\dagger$–0.696 | 0.057–0.069 |
| AA composition | 0.696–0.772 | 0.072–0.088 |
| TSS (DNABERT-2) | 0.217–0.276$^\dagger$ | −0.013––0.010 |

#### C.2 Additional robustness analyses

Table C2: TSS-window family macro-F1 on the homology-aware split and on a genomic-interval-disjoint split, in which no window overlaps a window in another partition. On the homology split, 212 of the 459 scored test genes (46.2%) share window sequence with a training or validation gene. Encoders at their validation-selected whole-window pooling rule.

| Source | Homology F1 | Disjoint F1 | $\Delta$ |
|---|---:|---:|---:|
| Shuffled labels (median of 200) | — | 0.167$^\dagger$ | — |
| TSS 4-mer | 0.283 | 0.281 | -0.002 |
| DNABERT-2 | 0.361 | 0.258 | -0.103 |
| NT-v2 | 0.347 | 0.300 | -0.047 |
| GENA-LM | 0.306 | 0.340 | +0.033 |
| HyenaDNA | 0.258 | 0.264 | +0.006 |
| Enformer (whole window) | 0.336 | 0.413 | +0.077 |
| Enformer (central 2,048 bp) | 0.402 | 0.411 | +0.009 |

Table C3: Paired cluster-bootstrap differences on the same test genes (1,000 resamples). The four primary tests carry Holm-adjusted one-sided $p$; every other block is exploratory with unadjusted $p$. Blocks: GenePT comparisons on CDS; CDS against the TSS window per encoder (disjoint split); Ends + Mean against Mean repeated three times at three times its $C$, the matched-regularisation control; the selected encoder against composition plus log CDS length; and the primary tests with each label-noise set masked from scoring (gene counts are the test genes removed).

| Comparison | Δ [95% CI] | $p$ |
|---|---|---|
| **Primary, macro-F1 (Holm-adjusted $p$)** | | |
| NT-v2 (Ends + Mean + Max) − CDS 6-mer | +0.054 [-0.011, +0.126] | 0.108 |
| NT-v2 (Ends + Mean + Max) − AA 3-mer | -0.075 [-0.137, -0.001] | 0.980 |
| ESM-2 650M − NT-v2 (Ends + Mean + Max) | +0.261 [+0.200, +0.322] | 0.004 |
| NT-v2 (Mean (Boundary-including)) (CDS) − NT-v2 (Mean) (TSS) (disjoint split) | +0.393 [+0.312, +0.479] | 0.004 |
| **Exploratory, GenePT $R^2$ (unadjusted $p$)** | | |
| AA 3-mer − AA 2-mer | +0.021 [+0.013, +0.029] | 0.001 |
| AA 3-mer − CDS 6-mer | +0.042 [+0.031, +0.054] | 0.001 |
| DNABERT-2 (Mean (Boundary-including)) − CDS 6-mer | +0.026 [+0.018, +0.034] | 0.001 |
| DNABERT-2 (Mean (Boundary-including)) − AA 3-mer | -0.015 [-0.031, +0.000] | 0.971 |
| ESM-2 650M − AA 3-mer | +0.096 [+0.082, +0.111] | 0.001 |
| ESM-2 650M − DNABERT-2 (Mean (Boundary-including)) | +0.111 [+0.087, +0.134] | 0.001 |
| **CDS − TSS window, same genes, disjoint split, macro-F1 (unadjusted $p$)** | | |
| DNABERT-2 | +0.380 [+0.295, +0.452] | 0.001 |
| NT-v2 | +0.393 [+0.312, +0.479] | 0.001 |
| GENA-LM | +0.267† [+0.173, +0.371] | 0.001 |
| HyenaDNA | +0.388 [+0.307, +0.455] | 0.001 |
| 4-mer (control) | +0.268 [+0.195, +0.346] | 0.001 |
| **CDS − TSS window, same genes, disjoint split, GenePT $R^2$ (unadjusted $p$)** | | |
| DNABERT-2 | +0.060 [+0.046, +0.071] | 0.001 |
| NT-v2 | +0.038 [+0.027, +0.049] | 0.001 |
| GENA-LM | +0.047 [+0.035, +0.056] | 0.001 |
| HyenaDNA | +0.052 [+0.040, +0.063] | 0.001 |
| 4-mer (control) | +0.048 [+0.039, +0.058] | 0.001 |
| **Ends + Mean − Mean ×3 at $3 \times C$, macro-F1 (unadjusted $p$)** | | |
| DNABERT-2 | -0.021 [-0.062, +0.020] | 0.838 |
| NT-v2 | -0.001 [-0.026, +0.028] | 0.509 |
| GENA-LM | +0.086† [+0.017, +0.153] | 0.006 |
| HyenaDNA | -0.016 [-0.045, +0.014] | 0.862 |
| **Encoder − composition with log CDS length, macro-F1** | | |
| NT-v2 (Ends + Mean + Max) − CDS 6-mer + length | +0.051 [-0.017, +0.122] | 0.064 |
| NT-v2 (Ends + Mean + Max) − AA 3-mer + length | -0.089 [-0.153, -0.020] | 0.994 |
| **Without noisy TF labels (84 test genes, 79 disjoint), macro-F1** | | |
| NT-v2 (Ends + Mean + Max) − CDS 6-mer | +0.047 [-0.025, +0.119] | 0.100 |
| NT-v2 (Ends + Mean + Max) − AA 3-mer | -0.076 [-0.140, -0.006] | 0.984 |
| ESM-2 650M − NT-v2 (Ends + Mean + Max) | +0.255 [+0.192, +0.323] | 0.001 |
| NT-v2 (Mean (Boundary-including)) (CDS) − NT-v2 (Mean) (TSS) (disjoint split) | +0.420 [+0.331, +0.503] | 0.001 |
| **Without non-protein-kinase labels (23 test genes, 24 disjoint), macro-F1** | | |
| NT-v2 (Ends + Mean + Max) − CDS 6-mer | +0.054 [-0.020, +0.133] | 0.082 |
| NT-v2 (Ends + Mean + Max) − AA 3-mer | -0.066 [-0.133, +0.001] | 0.972 |
| ESM-2 650M − NT-v2 (Ends + Mean + Max) | +0.254 [+0.188, +0.316] | 0.001 |
| NT-v2 (Mean (Boundary-including)) (CDS) − NT-v2 (Mean) (TSS) (disjoint split) | +0.392 [+0.314, +0.473] | 0.001 |
| **Without templated summaries (55 test genes), GenePT $R^2$** | | |
| DNABERT-2 (Mean (Boundary-including)) − CDS 6-mer | +0.024 [+0.018, +0.031] | 0.001 |
| DNABERT-2 (Mean (Boundary-including)) − AA 3-mer | -0.022 [-0.032, -0.012] | 1.000 |
| ESM-2 650M − DNABERT-2 (Mean (Boundary-including)) | +0.119 [+0.104, +0.135] | 0.001 |

Table C4: Ridge-to-GenePT alignment under measures that do not depend on the GenePT basis. Pooled $R^2$ weights each dimension by its variance; retrieval@5 is the share of the 459 scored CDS test genes (465 for the TSS row, on the disjoint split) whose true GenePT vector ranks in the top five by cosine to the prediction (chance 1.1%); median rank of the true vector.

| Method | Macro-$R^2$ | Pooled-$R^2$ | Retr.@5 | Med. rank |
|---|---|---|---|---|
| ESM-2 650M (upper bound) | 0.176 | 0.187 | 27.5% | 18 |
| ESM-2 150M | 0.158 | 0.169 | 20.3% | 31 |
| DNABERT-2, Mean (Boundary-including) (best DNA) | 0.065 | 0.072 | 5.0% | 96 |
| NT-v2, Mean (Boundary-including) | 0.043 | 0.051 | 6.3% | 79 |
| GENA-LM, Ends + Mean | 0.049 | 0.056 | 5.9% | 91 |
| HyenaDNA, Ends + Mean | 0.029 | 0.035 | 5.2% | 111 |
| TSS DNABERT-2, Mean | -0.011 | -0.010 | 2.4% | 195 |
| Shuffled-label control (4-mer, median of 200) | -0.027 | -0.026 | 1.1% | 230 |

Table C5: TSS-window family macro-F1 under whole-window pooling (validation-selected rule) versus TSS-Anchored pooling, each with its 95% cluster-bootstrap CI; Enformer as whole-window mean versus central 2,048 bp readout. Δ: anchored minus whole window on the same test genes, paired bootstrap; an anchored cell is bold where that CI lies above 0. Chunk comp.: the validation-selected composition baseline (4-mer + GC or 6-mer) of the same chunk. Shuffled-label median: 0.167†.

| Model | Whole window | TSS-Anchored [95% CI] | Δ [95% CI] | Chunk comp. |
|---|---|---|---|---|
| **Homology-aware split** | | | | |
| TSS 4-mer | 0.283 [0.228, 0.331] | — | — | — |
| DNABERT-2 | 0.361 [0.276, 0.427] | 0.261 [0.214, 0.299] | -0.100 [-0.178, -0.010] | 0.293 |
| NT-v2 | 0.347 [0.284, 0.394] | 0.370 [0.311, 0.417] | +0.023 [-0.038, +0.095] | 0.304 |
| GENA-LM | 0.306 [0.241, 0.365] | 0.350 [0.285, 0.408] | +0.044 [-0.021, +0.112] | 0.291 |
| HyenaDNA | 0.258 [0.215, 0.296] | 0.298 [0.231, 0.340] | +0.040 [-0.032, +0.097] | 0.329 |
| Enformer | 0.336 [0.251, 0.393] | 0.402 [0.312, 0.460] | +0.067 [-0.021, +0.150] | — |
| **Genomic-interval-disjoint split** | | | | |
| TSS 4-mer | 0.281 [0.223, 0.331] | — | — | — |
| DNABERT-2 | 0.258 [0.199, 0.308] | 0.281 [0.235, 0.322] | +0.024 [-0.030, +0.075] | 0.301 |
| NT-v2 | 0.300 [0.239, 0.354] | 0.351 [0.285, 0.411] | +0.050 [-0.005, +0.106] | 0.336 |
| GENA-LM | 0.340 [0.266, 0.406] | 0.367 [0.304, 0.427] | +0.027 [-0.058, +0.120] | 0.262 |
| HyenaDNA | 0.264 [0.220, 0.305] | 0.276 [0.224, 0.330] | +0.012 [-0.055, +0.079] | 0.325 |
| Enformer | 0.413 [0.331, 0.487] | 0.411 [0.345, 0.472] | -0.001 [-0.047, +0.044] | — |

Table C6: Partition composition: genes, the share in singleton 40% protein clusters, the median cluster size, the share in clusters of 10 or more, and olfactory receptors among GPCRs. A cluster split with family quotas places the largest clusters in training, so test is mostly genes without close paralogs.

| Partition | Genes | Singleton | Median | In ≥10 | ORs / GPCRs |
|---|---|---|---|---|---|
| **Homology-aware split (CDS primary)** | | | | | |
| Train | 2,271 | 19% | 3 | 33% | 388 / 414 |
| Val | 486 | 87% | 1 | 0% | 7 / 88 |
| Test | 487 | 87% | 1 | 0% | 6 / 89 |
| **Genomic-interval-disjoint split (TSS primary)** | | | | | |
| Train | 2,271 | 29% | 3 | 31% | 346 / 414 |
| Val | 487 | 61% | 1 | 11% | 55 / 88 |
| Test | 486 | 62% | 1 | 0% | 0 / 89 |

##### C.3 TSS-window genomic-overlap audit

To confirm that the TSS windows probe a substrate distinct from CDS, we measured how much of each 196,608 bp window overlaps the target gene's own coding sequence, using the GRCh38 Ensembl release 115 gene annotation. Across all 3,244 windows the mean target-CDS fraction is 0.008 (∼1%). The rest is the gene's own introns (18%) and untranslated exons (1%), the exons (11%) and introns (44%) of neighbouring genes, and intergenic sequence (25%). Because so little of the gene's own coding sequence leaks back into the window, the substrate ablation is interpreted on the unmasked window, with no explicit coding mask applied.

##### C.4 Split leakage audit

To measure what a random split leaks, we counted the test genes whose 40% protein cluster also contains a train or validation gene. On the random split 271 of 487 test genes (56%) have such a relative; on the homology split, none do by construction. The score gap between the two splits is reported in §3.5 and Table B4.

#### D Code Reproducibility

The code is at `github.com/Austin-Senna/dna_to_text` under the MIT licence; data and results are CC BY 4.0, except files derived from NT-v2 outputs, which keep NT-v2's CC BY-NC-SA 4.0. The tag `mlcb2026-camera-ready` marks the code behind every number in this paper, with every dependency pinned. Embedding extraction needs a GPU; every later step runs on CPU. A single script refits every probe cell from a clean tree, and an independent minimal reimplementation (NumPy, pandas, scikit-learn) reproduces the headline cells from the stored features. Sequences, embeddings, chunk caches, stored test predictions, the gene table and the HGNC snapshot it was built from are deposited on Zenodo (10.5281/zenodo.23031890), so every step after extraction can be rerun from the deposit without a GPU.

##### D.1 Deterministic settings and hyperparameters

**Splits.** The primary homology-aware split: each gene's canonical CDS is translated and clustered with MMseqs2 `easy-cluster` at 40% identity and 80% bidirectional coverage (`-min-seq-id 0.4 -c 0.8`), and whole clusters are assigned to partitions by a greedy largest-cluster-first rule that keeps 70/15/15 family proportions (seed 42). The split holds 2,271 train, 486 validation and 487 test genes (1,751 clusters over 3,244 genes); seeds 1, 7 and 123 give the replicate splits. The TSS-disjoint split groups genes that share a protein cluster or an overlapping window. Before scoring, a validation or test gene is masked when an all-against-all MMseqs2 search on full-length proteins finds a partner in an earlier partition at $E \leq 10^{-3}$, ≥40% identity and ≥80% coverage of both sequences (on the disjoint split, also an overlapping window); training genes are never masked.

**Probes.** Features are standardised inside every fit, with the scaler fit on the rows being fit. Classification is scikit-learn logistic regression (`lbfgs`, `tol=1e-6`, `max_iter=5000`, no class weighting); regression onto GenePT is ridge regression on unscaled targets. $C$ and $\alpha$ start on the grid $\{10^{-4}, \ldots, 10^{4}\}$, which extends one decade at a time while the validation pick sits on its edge, up to $10^{-6}$ and $10^{6}$ for $C$ and $10^{9}$ for $\alpha$. Extension stops when a decade gains no more than $10^{-4}$ or a fit fails to converge. A few cells, none behind a primary test, selected a value adjacent to a non-converged fit and keep their validation pick. Each hyperparameter, pooling rule, composition $k$ (nucleotide 4 or 6; amino acid 1, 2 or 3) and encoder is chosen on validation (macro-F1, or macro-$R^2$ for GenePT), separately on every split; the probe is then refit on train+validation and scored once on test. Every fit runs on one BLAS thread. A refit of the whole protocol on 6 threads, and another on a different OpenBLAS kernel, changes 44 of 764 cells (19 select a different $C$; test macro-F1 moves by at most 0.042) and no primary test; those digits carry a dagger in the appendix.

**Statistics.** Statistics are computed from the stored test predictions without refitting. Intervals and one-sided paired $p$-values come from 1,000 bootstrap resamples (seed 42) of whole groups: 40% protein clusters for CDS cells on the homology split, window-and-protein groups for TSS cells and on the disjoint split, joined through every protein pair that passes the purge rule and, for GenePT, through genes that share a templated summary. Resampling is by group, not stratified by family. Holm correction covers the four primary tests; every other test is exploratory and unadjusted. Each null band is the central 95% of 200 refits with train and validation labels shuffled (seeds 0–199), each re-selecting the regularisation strength; test labels are never shuffled.

**Translation.** Amino-acid $k$-mers and ESM-2 read each CDS translated through internal stops (as X), dropping a terminal stop and any trailing 1–2 nucleotides; the clustering behind the splits translates to the first stop. In translation a codon with a non-ACGT base becomes X, and amino-acid $k$-mers containing X are skipped; the codon baseline skips such codons. Of the canonical CDS, 18 are irregular: 17 carry an internal stop (14 of them olfactory receptors) and 1 an ambiguous base.

**Extraction.** Models run frozen, in inference mode, at revisions pinned in the code. CDS chunks are 510 content tokens overlapping by 64 for DNABERT-2 and GENA-LM, 998 overlapping by 64 for NT-v2, and 8,192 overlapping by 512 for HyenaDNA, which has no boundary tokens. TSS windows are 196,608 bp in gene orientation, centred on the canonical transcript's $5'$ end (Ensembl release 115); windows that run past a chromosome end are padded with N (8 genes, 498 to 54,023 bp), the same input for every model. Enformer's whole-window summary averages all 896 output bins of 128 bp (the central 114,688 bp; the outer flanks are attention context only), and its central summary averages the central 16 bins (2,048 bp).

### E Changes Since Submission

The camera-ready reruns every probe under a corrected protocol. We fixed the four primary tests (§3) after we had seen, on the same test genes, the submitted results and early reruns of a few cells under corrected settings, and before the first full run of the corrected protocol. After that trial run, an audit changed how the bootstrap groups related genes (Appendix D.1); this moved the tests' intervals and Holm $p$-values but not their point estimates, and changed no test's outcome. The tests are therefore not pre-registered. The submitted version had the following problems; correcting them changed the numbers (Table E1):

- **Pooling was selected on test,** not on validation as the submitted Methods said (the headline cell, NT-v2 with Ends + Mean + Max, was such a pick), and DNABERT-2 was fixed as the TSS representative by an earlier test pick. Every choice is now made on validation.
- **GENA-LM ran untrained.** Under the library version we used, GENA-LM's pretrained weights silently failed to load, so every submitted GENA-LM number came from a randomly initialised network. The pipeline now checks the loaded parameters against the weights file.
- **HyenaDNA read an untrained token.** A CLS token HyenaDNA was never trained with sat at the start of every input, so every HyenaDNA rule read it, and its CLS feature was the same for every gene. HyenaDNA now reads raw sequence, and boundary-token rules are undefined for it.
- **TSS windows.** Windows of minus-strand genes were not reverse-complemented, and the outermost gene start stood in for the TSS. Windows are now in gene orientation, centred on the canonical transcript's $5'$ end.
- **Enformer.** Tables mixed its readouts; it is now reported as the mean of all 896 output bins (whole window) and of the central 16 (central readout).
- **Translation.** The 17 CDS with an internal stop were cut at it for the amino-acid baselines

and ESM-2; they now translate through it.

- **Probes.** Every probe was fit on unscaled features, logistic fits stopped at tolerance $10^{-4}$ or 2,000 iterations, results depended on the BLAS thread count, composition baselines were picked at the edge of the regularisation grid, and the nucleotide $k$ was fixed at 4. Features are now standardised inside every fit, grids extend past their edge, both $k$ are chosen on validation, and logistic fits run on one thread to tolerance $10^{-6}$ and up to 5,000 iterations (settings in Appendix D.1).
- **Evaluation.** The submission resampled genes individually, applied no multiple-comparison correction, and estimated chance from one label shuffle. Intervals now come from a cluster bootstrap with Holm adjustment over the four primary comparisons, chance is a band from 200 shuffles, test genes with a close homolog in an earlier partition are masked from scoring, and the TSS arm uses a split that also separates overlapping windows.

These submitted conclusions changed:

- the selected encoder's lead over nucleotide composition, which the submission called well clear, is not significant (Holm $p = 0.108$), and it does not reach amino-acid composition, which the submission called level with it;
- GENA-LM, which the submission placed below 4-mer composition, scores highest of the four encoders on test once its weights load (0.710 on CDS, 0.340 on the TSS window), though it is not the validation pick;
- the submission read the ranking of the rules that add the first and last chunks as positional structure; at matched regularisation they show an exploratory gain only for GENA-LM (Section 3.3);
- on GenePT, amino-acid 3-mer composition, above every DNA encoder in the submission, is no longer reliably above the selected encoder: the paired interval touches zero (Appendix Table C3);
- on the TSS window, the encoders no longer clear composition: the submitted table put all four above the 4-mer (0.244 to 0.326 vs. 0.181); they now sit around it (0.258 to 0.340 vs. 0.281);
- the TSS window sits above chance. The submission's single-shuffle reference (0.224) was high; the 200-shuffle median is 0.167[†]. The encoders' margin over chance is about what it was, now on the disjoint split (homology-split values in Table C2), and the window stays far below CDS on the same genes (§3.4.1).

Table E1: Submitted versus camera-ready values for the main quantities: the submitted value, this version's value, and the cause; full values are in Appendix B. CDS rows are on the homology-aware split except the random-split rows; TSS rows were submitted on the homology-aware split and are now on the disjoint split. Macro-F1 unless marked $R^2$. "Selected" is the submission's best-on-test cell and this version's validation pick. Two causes apply to every row and are not repeated: the corrected probe protocol (standardised features, extended grids, tighter convergence, one thread) and, except on the random split, masking test genes with a close homolog. Row causes: P, pooling and encoder selected on validation; K, composition $k$ chosen on validation; T, translation through internal stops; G, GENA-LM weights loaded; H, HyenaDNA's untrained CLS token removed; W, TSS windows re-oriented and re-centred; E, Enformer readouts separated; S, cluster bootstrap with Holm adjustment; C, chance from 200 shuffles; D, disjoint split; —, the two shared causes only.

| Quantity | Submitted | This version | Cause |
|---|---:|---:|---|
| *Family classification, CDS* | | | |
| Selected DNA encoder (NT-v2) | 0.727 | 0.693 | — |
| NT-v2 − nucleotide comp. | well clear | +0.054 [−0.011, +0.126] | K, S |
| NT-v2 − amino-acid comp. | level | −0.075 [−0.137, −0.001] | K, T, S |
| Nucleotide composition reference | 0.633 (4-mer) | 0.639 (6-mer) | K |
| Amino-acid composition reference | 0.735 (AA 2-mer) | 0.767 (AA 3-mer) | K, T |
| ESM-2 650M | 0.960 | 0.954 | T |
| DNABERT-2 | 0.654 | 0.628 | P |
| HyenaDNA | 0.641 | 0.633 | H, P |
| GENA-LM (submitted: untrained) | 0.415 | 0.710 | G, P |
| NT-v2, pooling range | 0.519–0.727 | 0.504–0.698 | — |
| Chance | 0.224 (one shuffle) | 0.167 (median of 200) | C |
| *GenePT regression, CDS ($R^2$)* | | | |
| Selected DNA encoder (DNABERT-2) | 0.077 | 0.065 | P |
| AA 3-mer | 0.090 | 0.080 | T |
| CDS 4-mer | 0.045 | 0.032 | — |
| ESM-2 650M | 0.181 | 0.176 | T |
| *Family classification, TSS window* | | | |
| Selected TSS encoder (DNABERT-2) | 0.326 | 0.258 | W, D, P |
| NT-v2 | 0.313 | 0.300 | W, D, P |
| GENA-LM (submitted: untrained) | 0.244 | 0.340 | G, W, D, P |
| TSS 4-mer | 0.181 | 0.281 | W, D |
| Enformer central readout | 0.401 | 0.411 | W, E, D |
| *Random split* | | | |
| NT-v2, CDS | 0.828 | 0.801 | P |
| AA 2-mer, CDS | 0.837 | 0.854 | T |
| NT-v2, TSS | 0.447 | 0.487 | W, P |
